# Biological control of ion transport, redox activity, and nucleation during biogenic synthesis of CdS nanoparticles

**DOI:** 10.64898/2026.04.02.716127

**Authors:** Nicolas Bruna, Fengjie Zhao, Damo Nair, Rina Okuda, James Q. Boedicker

**Affiliations:** Department of Physics and Astronomy, University of Southern California, Los Angeles, CA, 90089, USA; Department of Biological Sciences, University of Southern California, Los Angeles, CA, 90089, USA

**Keywords:** biogenic synthesis, quantum dots, synthetic biology, metal ion uptake, cadmium

## Abstract

Cells have the potential to utilize biological pathways to synthesize semiconductor nanomaterials, such as CdS quantum dots. As in chemical reaction schemes, biogenic synthesis requires control of the concentration and redox state of starting materials during the nucleation and growth of nanoparticles. Biological pathways regulate these key processes of particle synthesis, and manipulation of such pathways enables biological control of multiple aspects of nanoparticle synthesis. Here, strains of *Escherichia coli* were engineered to biosynthesize cadmium sulfide (CdS) quantum dots through the coordinated action of three pathways controlling sulfide generation, cadmium uptake, and nanoparticle nucleation. When exposed to low, micromolar concentrations of external cadmium, strains combining all three pathways produced CdS quantum dots. The synthesis of nanoparticles, nanoparticle yield, and nanoparticle size depended on the combination of pathways found in each strain. Cells lacking all three pathways produced no detectable nanomaterials, cells with specific combinations of one or two pathways produced small particles in the range of 1.95 to 7.9 nm, and cells with all three pathways produced the largest particles with average diameters of 11.78 nm. These results demonstrate that cells can be engineered to control multiple aspects of biogenic nanoparticle synthesis and that these pathways act together to tune the biosynthesis of semiconductor nanomaterials within cells.

**Importance:** Microbes synthesize materials, including metallic and semiconductor nanomaterials. This capability stems from the natural ability of microbes to interact with and precisely manipulate metal atoms. Here, multiple biological pathways were combined within a single strain of *Escherichia coli*, creating a cell capable of producing CdS nanoparticles. This engineered cell controls multiple steps of particle synthesis, including metal uptake, reduction of starting materials, and binding cadmium and sulfide ions to initiate particle formation. Metal uptake by the cells was improved through the modification of a metal ion transport protein, improving cadmium uptake across the outer membrane and creating higher concentrations of cadmium within the cell. Cells with all three pathways were able to produce CdS nanoparticles, called quantum dots, even when exposed to low concentrations of external cadmium. This biotechnology enables nanomaterial synthesis under environmentally friendly conditions and may improve technologies using bacteria to clean up toxic metals.

## Introduction

Many biomolecules have evolved to interact with a variety of atoms and small molecules, including metal ions. Metal-handling pathways within cells are often associated with integrating metals ions into enzymes or mitigating metal toxicity, however prior work has repurposed these pathways for the synthesis of nanoscale materials (Sharma et al., 2019; Narayanan & Sakthivel, 2010; Dundas et al., 2018; Era et al., 2022; Contreras et al., 2025), including inorganic materials such as semiconductor quantum dots (Kang et al., 2008, Lei et al., 2025; Shafiei et al., 2025; Vargas-Reyes et al., 2024). Such biogenic material synthesis within living cells involves the uptake and processing of raw materials from the environment, followed by the nucleation and growth of the material (Naughton et al., 2021). Genetic engineering offers the possibility to combine multiple pathways within a cell to regulate material synthesis, adjusting the conditions for material synthesis and tuning the properties of the resultant biogenic materials.

Here biological pathways for metal uptake, sulfur reduction, and particle nucleation are combined to control the biogenic synthesis of CdS quantum dots within living cells of *Escherichia coli*. Semiconductor quantum dots are nanoscale fluorescent particles that exhibit size-dependent electronic and optical properties as a consequence of quantum confinement, enabling applications in optoelectronics, sensing, and bioimaging (Le & Kim, 2023; Michalet et al., 2005; Alivisatos, 1996). A wide variety of quantum dots have been developed, including classical II–VI materials such as CdS, CdSe, and CdTe, each offering tunable optical and electronic characteristics (Houtepen et al., 2025). Traditionally, these nanocrystals are synthesized through high-temperature chemical routes that rely on reactive organometallic precursors, hazardous solvents, and substantial energy input, raising concerns about environmental burden and process scalability (Yu et al., 2003; Qu, L. & Peng, X., 2002; Peng, Z.A. & Peng, X., 2001; Peng X. et al., 2000).

In contrast to chemical synthesis, biological systems enable quantum dots formation through cellular metabolism, producing fluorescent nanoparticles under biologically compatible conditions. Organisms such as bacteria, fungi, yeasts, microalgae, and plant extracts have been employed for biosynthesis, enabling environmentally friendly production of diverse quantum dots including CdS, CdSe, CdTe, and Ag_2_Se (Gangan et al., 2023; Liu et al., 2021; Bruna et al., 2019; Zhang et al., 2018; Shivaji et al., 2018; Chen et al., 2014; Syed et al., 2013). These biologically produced nanoparticles have demonstrated applications in antimicrobial treatments, bioimaging, and solar energy devices (Shivaji et al., 2018; Órdenes-Aenishanslins et al., 2014).

Beyond native metabolic capabilities, genetic engineering has emerged as a powerful strategy to enhance and control the biological synthesis of semiconductor quantum dots (Edmundson et al., 2014). Early studies demonstrated that recombinant *E. coli* can be genetically engineered to biosynthesize CdS nanocrystals through the introduction of metal-binding or sulfur-related genetic elements, establishing proof of principle for genetically controlled nanoparticle formation (Kang et al., 2008). Subsequent work revealed that increasing intracellular thiol pools via overexpression of glutathione biosynthesis enzymes enhances the biosynthesis of both CdS and CdTe quantum dots in engineered *E. coli*, highlighting the central role of redox balance, regulated intracellular cadmium handling, and detoxification pathways in quantum dot formation and fluorescence properties (Chen et al., 2009; Monrás et al., 2012). Complementary approaches introduced plasmid-encoded CdS-binding peptides to promote nucleation and control particle growth, enabling the production of fluorescent CdS quantum dots with improved optical properties in engineered *E. coli* (Mi et al., 2011). More recently, recombinant expression of metallothioneins has been employed to enhance intracellular cadmium binding and retention, thereby facilitating the intracellular assembly of CdSe quantum dots in both *E. coli* and non-model bacteria, such as the photosynthetic *Rhodopseudomonas palustris* (Jia et al., 2023; Zhou et al., 2022). These studies collectively illustrate that targeted genetic interventions can modulate metal uptake, intracellular cadmium availability, and nucleation dynamics, thereby shaping quantum dot size, composition, and optical behavior.

In addition to intracellular metabolism, metal uptake itself has emerged as a critical and engineerable determinant of biogenic nanomaterial synthesis. Building on these advances, our recent work demonstrated that regulating a divalent metal ion transporter provides an additional level of control over biogenic nanoparticle synthesis. In Gangan et al. (2023), expression of a metal ion transporter was used to regulate intracellular cadmium availability, controlling nanoparticle formation. This work established metal transport as a limiting step that can be genetically modulated independently of downstream biosynthetic pathways, providing a modular strategy to couple metal uptake with intracellular nucleation processes.

In this study, three biochemical pathways were combined to control the synthesis of CdS quantum dots, integrating pathways to control sulfur metabolism, metal uptake, and nucleation to drive CdS quantum dot biosynthesis in *E. coli*. Specifically, sulfide availability was regulated through induction of the *phsABC* operon, enabling *E. coli* to reduce sodium thiosulfate and create an intracellular sulfur source for nanoparticle formation. Intracellular cadmium levels were enhanced via expression of the divalent metal transporter ZupT targeted to the outer membrane (ZupT_OM), and the impact of this transport system on Cd incorporation was quantified using a GFP-based cadmium biosensor. Expression of a nucleating peptide known at A7 promoted nanoparticle nucleation, facilitating CdS quantum dot formation. The combination of these pathways controlled under which conditions quantum dots were synthesized and the size of the resulting material. This work demonstrates how coordinated metabolic and genetic engineering can enable efficient, tunable, and environmentally compatible biosynthesis of CdS quantum dots, while providing quantitative insight into the intracellular metal dynamics that govern nanoparticle formation.

## Results

*E. coli* cells were genetically engineered to control the synthesis of chalcogenide nanoparticles within living bacteria. Nanoparticle synthesis within cells relies on three key processes: the import of the reactants for material synthesis, redox reactions with these starting materials, and nanoparticle nucleation (Naughton et al., 2021). Here, each of these processes was biologically regulated by genetic modification of the cells. Cadmium ions and thiosulfate were added to cultures of these engineered cells as the starting materials for synthesis CdS nanoparticles.

### Controlling sulfide production

To control the processing of starting materials using redox enzymes, a *Salmonella enterica phsABC* operon encoding the thiosulfate reductase pathway under the control of an IPTG-inducible promoter was integrated into the genome of *E. coli BL21(DE3)* using the markerless guide-RNA-assisted targeting system INTEGRATE (Vo et al., 2021). The *phsABC* operon was integrated into the chromosome of *E. coli BL21(DE3)* generating strain *E. coli::phsABC*. Gibson assembly was used to modify the INTEGRATE plasmid pSpin with a guide RNA and the *phsABC* cargo. The construct expressing *phsABC* contained a promoter inducible by IPTG. DNA gels of PCR products confirmed insertion of this construct into the genome at the target site, see Figure S1.

The *phsABC* pathway encodes for membrane proteins capable of converting thiosulfate to sulfide, as shown in Figure 1A. A sulfide assay was used to confirm the functionality of the *phsABC* pathway in *E. coli::phsABC*. Lead acetate paper, which turns black in the presence of H_2_S, was placed in the head space above 3 mL cultures of cells with 100 µM thiosulfate added (Liu et al., 2022). As shown in Figure 1B, the lead acetate paper turned black for a culture of *E. coli::phsABC* with 1 mM of IPTG added to induce expression of *phsABC*. Controls with medium only, *E. coli BL21(DE3)* cells not containing the *phsABC* pathway, and *E. coli::phsABC* without addition of the inducer did not produce a dark color on the lead acetate paper. Additional controls for the H_2_S assay are shown in Figure S2. These results indicate strain *E. coli::phsABC* converted thiosulfate into detectable levels of H_2_S.

**Figure 1:**
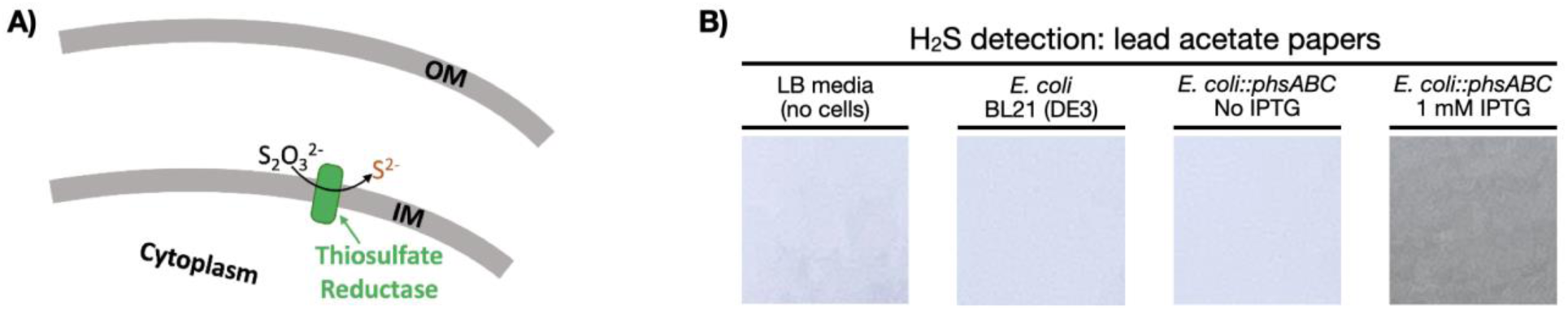
Thiosulfate reduction and H_2_S production in *E. coli::phsABC* strain. **(A)** Schematic representation of thiosulfate reduction by *E. coli::phsABC* strain. **(B)** Lead acetate paper assay for detection of H_2_S production after incubation in LB medium containing 100 µM thiosulfate under the following conditions: medium only (no cells), wild-type *E. coli BL21(DE3)*, *E. coli::phsABC* without IPTG, and *E. coli::phsABC* induced with 1 mM IPTG.

### Controlling cadmium ion uptake

Next, *E. coli* was modified to increase the uptake of external Cd ions. A native pathway in *E. coli* that contributes to the uptake of divalent metal ions, including Cd, is the ZupT metal permease (Taudte & Grass, 2010). The protein is expressed in the inner membrane of *E. coli*, but prior work has shown that fusion of a signal peptide to the N-terminus of *Z*upT resulted in localization of this permease to the outer membrane of *E. coli* and increased the uptake of zinc (Gangan et al., 2023). The strain carrying the plasmid expressing the outer membrane targeted *Z*upT is designated *E. coli* ZupT_OM, resulting in expression of ZupT on both the inner and outer membranes as depicted in Figure 2A. To more thoroughly characterize Cd uptake in this strain, Cd concentrations in the cytoplasm were measured using a Cd-responsive transcription factor (Beabout et al., 2021). This technique was used to probe Cd uptake for strains with different combinations and variants of ZupT exposed to external Cd concentration from 1 - 25 µM.

**Figure 2.**
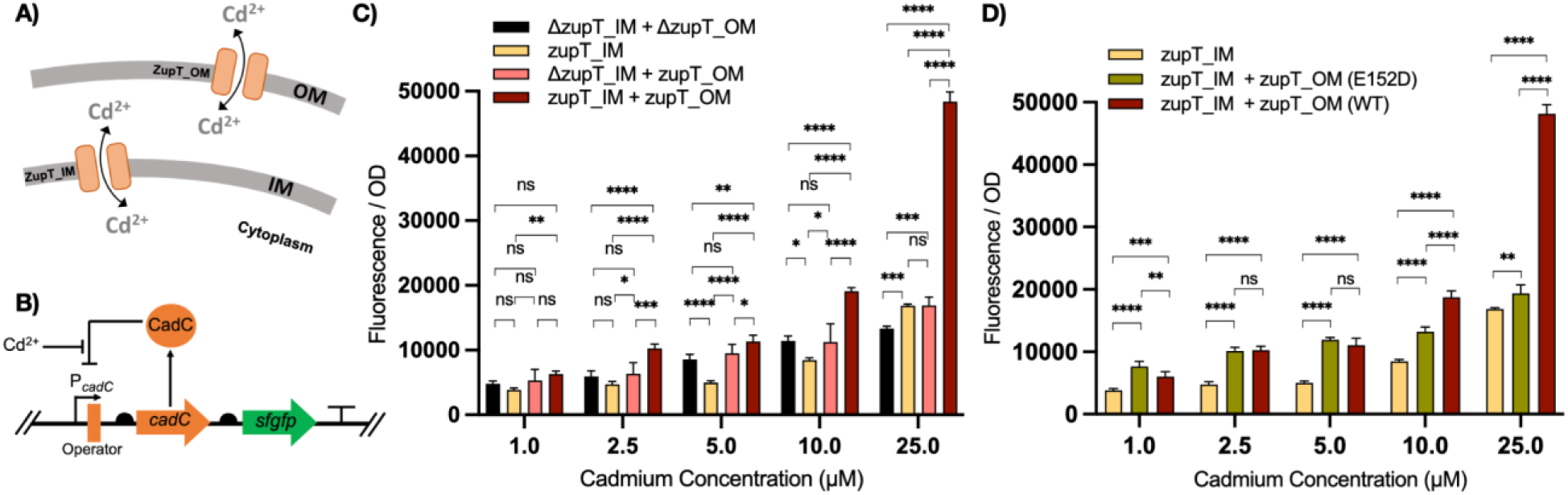
Cadmium uptake analysis in *E. coli* strains expressing ZupT in the outer membrane. **(A)** Schematic representation of cadmium transport across the outer membrane (OM) and inner membrane (IM). **(B)** Diagram of the cadmium biosensor reporter, which expresses GFP in response to the concentration of cytoplasmic cadmium. **(C)** Fluorescence intensity measured using the GFP-based reporter biosensor. Strains expressing different combinations of the *zupT* gene were compared: *E. coli ΔzupT_IM + ΔzupT_OM*, lacking *Z*upT in both the inner and outer membranes; *E. coli zupT_IM*, expressing *Z*upT only in the inner membrane; *E. coli ΔzupT_IM + zupT_OM*, expressing *Z*upT only in the outer membrane; *E. coli zupT_IM + zupT_OM*, expressing *Z*upT in both membranes. **(D)** Fluorescence intensity measured by the reporter biosensor in *E. coli zupT_IM*, *E. coli zupT_IM + zupT_OM (E152D)*, and *E. coli zupT_IM + zupT_OM (WT)*. The E152D mutation in *Z*upT is known to reduce cadmium uptake. Bars represent the mean of five measurements, and error bars indicate the standard deviation. Statistical significance is indicated as follows: ns, not significant; \**p* < 0.05; \*\**p* < 0.01; \*\*\**p* < 0.001; \*\*\*\**p* < 0.0001. All strains in C and D contain the plasmid pCad2.

The cadmium reporter construct is encoded on a plasmid that contains the CadC cadmium-responsive transcription factor from *Staphylococcus aureus* (Endo & Silver, 1995), as shown in Figure 2B. In the absence of cadmium, CadC binds to the promoter of the *gfp* reporter gene to repress GFP expression. When there is sufficient Cd in the cytoplasm of cells, Cd binds to CadC to derepress the expression of GFP. To use this Cd assay, strains were transformed with the pCad2 reporter plasmid. After growing cultures to OD₆₀₀ 0.2, variable concentrations of cadmium were added to the LB media for 3 hours. As shown in Figure S3, *E. coli BL21(DE3)* containing the reporter plasmid pCad2 had detectable levels of GFP fluorescence when more than 16 nM of Cd were added to LB media.

To demonstrate the ability of the outer membrane ZupT to impact cytoplasmic concentrations of cadmium, *E. coli* strains with different combinations of the ZupT protein were compared. *E. coli ΔzupT_IM + ΔzupT_OM is a* strain lacking both the inner and outer membrane versions of ZupT. *E. coli zupT_IM* represents *E. coli* with only the native inner membrane ZupT. Transforming the plasmid pZupT_OM expressing the outer membrane ZupT into *E. coli ΔzupT_IM* created strain *E. coli ΔzupT_IM + zupT_OM* with only an outer membrane ZupT. Transforming the same plasmid into *E. coli* zupT_IM resulted in a strain with both inner and outer membrane ZupT called *E. coli zupT_IM + zupT_OM*. The response of these strains to external concentrations of Cd between 1 and 25 µM is shown in Figure 2C.

At low concentrations of Cd (1-10 μM), the fluorescence of *E. coli zupT_IM* was similar to or less than that of *E. coli ΔzupT_IM + ΔzupT_OM*. However, at 25 µM external Cd, the fluorescence of *E. coli zupT_IM* was higher than that of *ΔzupT_IM + ΔzupT_OM*. This indicates the native inner membrane ZupT increases Cd uptake only at higher concentrations of Cd. *E. coli zupT_IM + zupT_OM* exhibited the highest fluorescence among all strains tested at all Cd concentrations, particularly at 10 and 25 μM, suggesting that the presence of ZupT in both the inner and outer membranes significantly increases Cd uptake into the cytoplasm. Together, these results indicate that at low micromolar external Cd concentrations, Cd does not efficiently cross to the cytoplasm with inner membrane ZupT alone. Adding the divalent metal ion permease ZupT to the outer membrane increases Cd uptake under these conditions. The inner membrane ZupT increased uptake only at 25 µM external cadmium, whereas the presence of both inner and outer membrane ZupT increased cadmium uptake even at 1 µM.

To further verify the ability of ZupT with the outer membrane translocation tag to act as a metal ion permease when expressed in the outer membrane, mutational analysis was performed. Prior work identified a non-synonymous mutation in ZupT that reduced cadmium transport, the *Z*upT E152D mutant (Taudte, N., & Grass, G., 2010). Residue 152 is part of the pore of the permease, through which metal ions translocate across the membrane. The E152D mutation was incorporated into the zupT_OM sequence using site directed mutagenesis to create pzupT_OM(E152D), see Figure S4. As shown in Figure 2D, strains expressing the mutant *zupT_OM* (E152D) had significantly lower fluorescence at both 10 and 25 µM cadmium. This point mutation impacting cadmium uptake supports that ZupT located in the outer membrane indeed acts as a metal ion permease.

### Nanoparticle synthesis in strains with engineered uptake, redox, and nucleation pathways

Given that the thiosulfate reductase and the metal ion permease control intracellular sulfide and Cd concentrations, an additional nanoparticle nucleation pathway was introduced by expressing CdS nucleation-associated peptides in the cytoplasm. The peptide named A7 (CNNPMHQNC) (Mao et al., 2004) was attached to the C-terminus of LacZ to increase CdS nucleation within the cytoplasm. The A7-lacZ fusion was expressed from plasmid pA7peptide using aTc induction (see Table S1).

Nanoparticle synthesis in engineered strains expressing these pathways (Figure 3a) was characterized by photoluminescence and absorbance measurements following cell collection and lysis. For synthesis, cells were grown with induction to an OD₆₀₀ of 0.2–0.3, followed by the addition of 100 µM sodium thiosulfate and 1 µM cadmium chloride.

**Figure 3:**
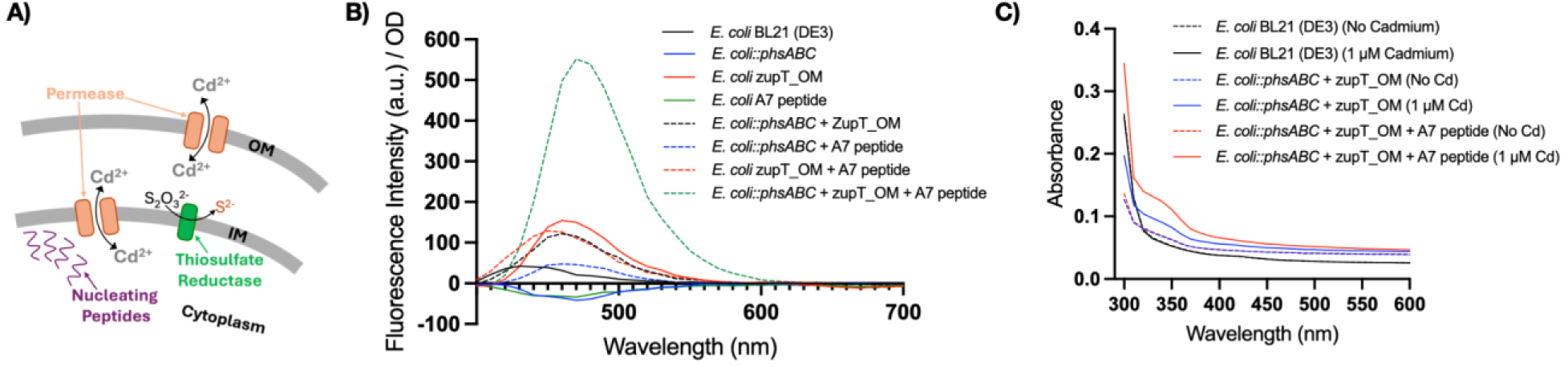
CdS nanoparticle synthesis by engineered strains of *E. coli*. (**A)** The strains contained pathways related to particle synthesis: the outer membrane permease ZupT, thiosulfate reductase PhsABC, and nucleation peptide A7. **(B)** Photoluminescence and **(C)** absorbance of nanoparticles harvested from cultures of cells expressing different combinations of the synthesis pathways. In B) 1 µM Cd was added to cultures during synthesis and in C) either 0 or 1 µM Cd was added to cultures during synthesis.

Cells were harvested by centrifugation, lysed, and the supernatant collected. The presence of CdS nanoparticles was quantified by detecting the characteristic photolumiscence emission of CdS nanoparticles around 470 nm when excited at 365 nm. The supernatant of the cell culture did not exhibit detectable photoluminescence, suggesting that the majority of nanoparticles collected were synthesized internally.

Photoluminescence of nanoparticles synthesized in strains with different combinations of pathways was measured, as shown in Figure 3b. These strains contained individual pathways (phsABC, zupT_OM, or A7 peptide), combinations of two pathways, or all three pathways. The host strain expressed ZupT_IM in all cases.

The sample from the WT strain *E. coli BL21(DE3)* had a low fluorescence, suggesting no nanoparticle synthesis. This low fluorescence is likely due to autofluorescence of cellular debris, such as autofluorescent proteins and NADPH which exhibit low fluorescence emission in the range of 400-500 nm (Croce & Bottiroli, 2014). The nanoparticles synthesized in strain *E. coli::phsABC* + zupT_OM + A7 peptide with all three pathways, dashed blue lines, had the highest peak in fluorescence near 470 nm. Nanoparticles synthesized by three additional strains, *E. coli zupT_OM*, *E. coli zupT_OM + A7 peptide*, and *E. coli::phsABC + zupT_OM*, had weaker fluorescence than *E. coli::phsABC + zupT_OM + A7 peptide* but higher than the WT strain. All the strains with detectable nanoparticles expressed zupT_OM, showing the importance of increasing cadmium uptake for internal nanoparticle synthesis. The nucleation peptide and thiosulfate reductase assisted in nanoparticle synthesis, and the combination of all three pathways greatly increased nanoparticle synthesis. Samples isolated from strains with only A7 peptide and PhsABC did not show detectable fluorescence, which suggests no nanoparticle synthesis. Samples isolated from strains *E. coli::phsABC + A7 peptide* showed low fluorescence similar to WT, but with a peak near 470 nm, which may indicate a low yield of nanoparticles.

Absorbance data shown in Figure 3C, confirms these conclusions. Samples containing CdS nanoparticles should have a shoulder, characteristic of CdS nanoparticles, in the absorbance spectrum near 350 nm (Lakowicz et al., 1999). Sample from *E. coli BL21(DE3)* cells and also all samples from cells without adding cadmium in the culture showed no shoulder. Sample from strain *E. coli::phsABC + zupT_OM* showed a small shoulder and sample from strain *E. coli::phsABC + zupT_OM + A7 peptide* showed a more pronounced shoulder and the highest absorbance, when cadmium was added to the cell culture.

### Nanoparticle size depends on the combination of pathways used for synthesis

To examine the properties of the synthesized nanoparticles, dynamic light scattering was used to measure particle size. Harvested particles synthesized following the protocol used in Figure 3 were measured in a Wyatt ZetaStar/Mobius. The sample from *E. coli BL21(DE3)* strain, which in Figure 3B showed a very low background fluorescence, had no detectable nanoparticles by light scattering, as shown in Figure 4. Strains *E. coli::phsABC* and *E. coli A7 peptide* also had no detectable nanoparticles, supporting the interpretation in Fig. 3B that these strains did not make nanoparticles. Cultures of other strains synthesized nanoparticles with average diameters between 2 and 12 nm. Strain *E. coli::phsABC + A7 peptide* showed nanoparticles with the smallest average diameter of 1.95 nm, suggesting that a lack of the ZupT_OM led to reduced uptake of Cd and therefore synthesis at a lower Cd concentration. Synthesis in strains with lower Cd concentrations would have delayed particle nucleation and slowed nanoparticle growth, resulting in smaller particles. Strain *E. coli::phsABC + zupT_OM + A7 peptide* produced the largest nanoparticles, with an average diameter of 11.78 nm (see Figure S5 for statistical testing). The synthesis of large nanoparticles suggests that strains with all three pathways created a chemical environment inside the cell amenable to nucleation and growth of CdS nanoparticles.

**Figure 4.**
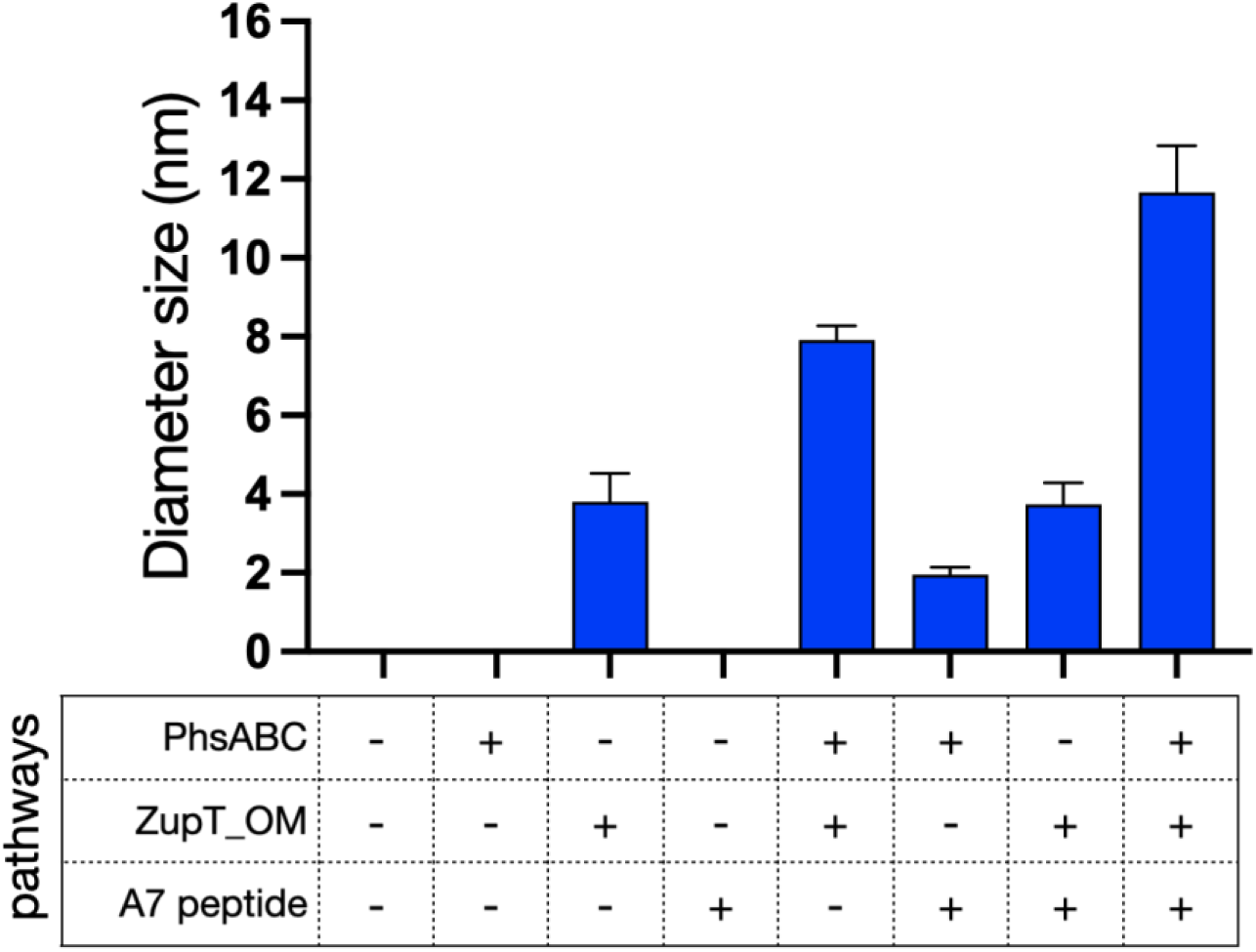
Dynamic light scattering analysis. Hydrodynamic diameter of CdS quantum dots synthesized by the bacterial strains containing the synthesis pathways indicated on the x-axis, as determined by light scattering. The y-axis shows the mean particle diameter in nanometers (nm), and error bars represent the standard deviation of four independent measurements (n = 4). No bar represents no nanoparticles detected.

## Discussion

CdS nanoparticles were synthesized using engineered strains of *E. coli* to manipulate multiple steps of particle synthesis. Genetic constructs were added to *E. coli* to influence cadmium uptake across the outer membrane, reduce thiosulfate to the sulfide, and assist with nanoparticle nucleation within the cytoplasm. Strains containing combinations of these pathways were able to synthesize CdS nanoparticles even at low external concentrations of Cd. Prior work synthesized CdS nanoparticles at external concentrations of Cd greater than 100 µM (Gu et al., 2025; Li et al., 2025; Vargas-Reyes et al., 2024; Yan et al., 2017; Chen et al., 2014; Chen et al., 2009). At such high concentrations of Cd, pathways to increase Cd uptake or assist with particle nucleation were not required, as high concentrations of Cd were likely found both extracellularly and intracellularly resulting in widespread and poorly controlled particle synthesis. The approach here was to use biological pathways to regulate multiple steps of particle synthesis to achieve nanoparticle synthesis at low concentrations of external Cd, at which a WT strain was unable to synthesize nanoparticles. As shown in Figure 3 and 4, some strains containing combinations of these pathways synthesized detectable levels of nanoparticles, and the strain containing all three pathways had the highest particle yield and the largest particles.

Particle synthesis results suggest that transport of cadmium across the outer membrane was an important barrier to nanoparticle formation. As shown in Figure 3B, the four strains with photoluminescence above the wildtype strain all contained ZupT_OM (*E. coli zupT_OM*, *E. coli::phsABC + zupT_OM*, *E. coli zupT_OM + A7 peptide*, and *E. coli::phsABC + zupT_OM + A7 peptide*). This suggests that when external cadmium was at 1 µM, a lack of intracellular cadmium prevented nanoparticle synthesis. The results of Figure 2 support this conclusion, as the addition of ZupT_OM to strains resulted in higher concentrations of cytoplasmic cadmium as compared to WT cells with ZupT_IM only. Prior work had shown that a strain of *E. coli* expressing *zupT_OM* increased the synthesis of CdS nanoparticles (Gangan et al 2023). Our Cd uptake results using the Cd reporter more directly show the impact of ZupT on cytoplasmic Cd concentrations. At external Cd concentrations of 5 and 10 µM, strain *E. coli zupT_IM* had a lower fluorescence than strain *E. coli ΔzupT_IM + ΔzupT_OM*. This may be due to zupT_IM assisting in the efflux of cytoplasmic Cd in these contexts. ZupT facilitates the passive transport of ions, enabling diffusion across the membrane in either direction. Potentially active transport, through other metal ion transport proteins on the inner membrane (Campoy et al., 2002; Patzer & Hantke, 1998), resulted in a reverse gradient of Cd across the inner membrane. In this case the presence of ZupT_IM would reduce cytoplasmic concentrations of Cd.

Work shown here also gives additional support for ZupT to function as a permease when expressed in the outer membrane. Mutation of a residue in the permease pore, known to reduce cadmium uptake, resulted in lower cytoplasmic cadmium concentrations as compared with the strain *E. coli zupT_IM + zupT_OM* (Figure 2D). This result suggests that ZupT expressed into the outer membrane contains a pore to facilitate the diffusion of cadmium through the membrane, as opposed to the expression of the protein in the outer membrane causing a general disruption of membrane function. These results suggest that the outer membrane is an important barrier to uptake Cd at low external concentrations (<25 µM). Overcoming this barrier, through strategies such as the outer membrane permease used here, may improve efforts to engineer strains for metal uptake and synthesis of nanomaterials.

This study also highlights the ability to biologically control the properties of biogenic nanomaterials. Whether or not detectable levels of nanomaterials were formed depended on the presence of ZupT_OM, and the size of the particles was dependent on which pathways were present within each strain. Adding the A7 peptide to the strain with PhsABC and ZupT_OM resulted in a 60% increase in particle diameter. This size increase is likely due to the increase in the rate of nanoparticle nucleation, allowing nanoparticles to form earlier in the process and grow to a larger size. The larger size of the particles is supported by photoluminescence data. Quantum dot size is related to the wavelength of light emitted by the particles, with larger particles emitted longer wavelengths. Particles made by strain *E. coli:phsABC + zupT_OM + A7 peptide* had a peak emission of 470 nm whereas particles made by strain *E. coli:phsABC + zupT_OM* had a peak emission of 450 nm. The absorbance shoulder shifting to a longer wavelength in Figure 3C also supports this larger size. Prior work has shown that properties of biogenic nanoparticles, such as particle size, depended on the time of synthesis (Gu et al., 2025; Darroudi et al., 2011; Bai et al., 2009). Here we demonstrate that the expression of proteins involved in nanoparticle synthesis can also control the properties of the synthesized nanoparticles.

Biogenic synthesis of nanoparticles is dependent on multiple pathways within the cell. The ability to genetically tune nanoparticle formation and size offers opportunities for applications in bioimaging, sensing, and optoelectronics. Furthermore, the modular design of these pathways provides a platform to extend this approach to other nanomaterials or host organisms. Finally, these findings establish a foundation for mechanistic studies into intracellular metal transport, redox chemistry, and nucleation, which can guide future efforts to optimize biogenic nanoparticle synthesis.

## Materials and Methods

### Bacterial strains, plasmids, and genetic modifications

*E. coli BW25113* was used to examine Cd uptake in strains expressing combinations of ZupT_OM and ZupT_IM. The Δ*zupT* mutant from the Keio collection was derived from host strain E. *coli BW25113* (Baba et al., 2006).

Other strains were derived from strain *Escherichia coli BL21(DE3)*. To enable thiosulfate-dependent H₂S production, the *phsABC* operon was integrated into the chromosome via the RNA-guided DNA insertion system INTEGRATE, which directs transposon insertion to a target site adjacent to a protospacer adjacent motif (PAM) (Vo et al., 2021; Jiang et al., 2015; Reisch & Prather, 2015). The *phsABC* operon from *Salmonella enterica*, encoding a thiosulfate reductase pathway, was cloned from plasmid pSB74 into the INTEGRATE plasmid pSpin together with the guide RNA using Gibson assembly, generating plasmid pSpin-*phsABC*. The resulting plasmid was transformed into *E. coli BL21(DE3)*, where the *phsABC* operon was integrated into the chromosome approximately 50 bp downstream of the target site at 3971363 (Park et al., 2020), generating strain *E. coli::phsABC*. The chromosomal *phsABC* operon is under IPTG-inducible control, and expression was induced with 1 mM IPTG. To enhance uptake of divalent metal ions and promote intracellular CdS formation, a subset of *E. coli* BL21 WT + *phsABC* strains were transformed with a pBAD24-derived plasmid expressing the outer membrane protein ZupT fused to an OmpA signal peptide (zupT_OM). Expression from this plasmid was induced with 1 mM arabinose. The nucleating peptide CNNPMHQNC was expressed from a pDSG372-derived plasmid as a LacZ fusion, resulting in cytoplasmic expression of the peptide. Expression from this plasmid was induced with 100 ng/mL anhydrotetracycline (aTc).

Plasmids used in this study are listed in Table S1. All constructs were generated using Gibson assembly or site-directed mutagenesis and verified by Sanger sequencing. Plasmids were transformed into competent cells by electroporation at 1.4 kV using an Eppendorf Eporator (Eppendorf, Germany). Strains were grown in LB medium supplemented with antibiotics as required for plasmid maintenance (100 µg/mL ampicillin and 50 µg/mL kanamycin). pCad2 is a modified version of the Cd-biosensor plasmid pCad1, (Beabout et al., 2021). Gibson assembly was used to replace the ColE1 origin of replication with a p15A origin and the kanamycin resistance marker with a chloramphenicol resistance cassette to allow selection in the Keio Δ*zupT* strain. pZupT_OM(E152D) is a variant of pzupT_OM, with a point mutation introduced in *zupT_OM* using site-directed mutagenesis. The E152D mutation in *zupT* reduces Cd uptake kinetics (Taudte & Grass, 2010). The presence of the E152D mutation was confirmed by Sanger sequencing, and the resulting decrease in fluorescence compared to the zupT_IM + zupT_OM(WT) strain confirmed the functional impairment of cadmium transport (Figure 2D).

### H₂S production through thiosulfate reduction

Hydrogen sulfide production was assessed in cultures of *E. coli* WT, *E. coli*::*phsABC*, *E. coli*::*phsABC* + zupT_OM, *E. coli*:: *phsABC* + zupT_OM + A7 peptide. Each strain was inoculated to an initial optical density of 0.05 with sodium thiosulfate added at a final concentration of 100 µM. Cultures were grown overnight in LB medium at 37 °C with shaking. H₂S release was monitored qualitatively using lead acetate papers (Lead Acetate Strips, Fisher Scientific, catalog number NC9506930), a standard method for detecting volatile sulfide species in bacterial cultures (Siegel, 1965; Shatalin et al., 2011; Zhu & Chu, 2022). Formation of dark lead sulfide deposits was recorded as evidence of H₂S production. Control cultures lacking thiosulfate were included to determine background sulfide levels. Pictures were taken using an iPhone 13 (Apple Inc., USA) equipped with the ProCam application.

### Analysis of ZupT-mediated cadmium uptake

To assess the contribution of ZupT to cadmium uptake in *E. coli*, the biosensor plasmid pCad2 was used, which expresses GFP in response to intracellular cadmium (Beabout et al., 2021). The pCad2 plasmid was transformed into four bacterial strains: *E. coli ΔzupT_IM + ΔzupT_OM*, *E. coli zupT_IM*, *E. coli ΔzupT_IM + zupT_OM*, and *E. coli zupT_IM + zupT_OM*. Cultures were incubated with increasing concentrations of cadmium chloride for 3 hours at 37 °C with shaking (Figure 2C). GFP fluorescence was measured as a proxy for intracellular cadmium levels using a TECAN Infinite M200 Pro plate reader, with excitation at 485 nm and emission collected at 515 nm. Fluorescence measurements were normalized to cell density by dividing by absorbance at 600 nm, and the background fluorescence per cell of the same strain without added Cd was subtracted.

### Biosynthesis of CdS Quantum Dots

Eight bacterial strains were used for CdS biosynthesis: *E. coli* WT; *E. coli::phsABC*; *E. coli::phsABC* + zupT_OM; *E. coli::phsABC* + zupT_OM + A7 peptide; *E. coli* zupT_OM; *E. coli* A7 peptide; *E. coli*::*phsABC* + A7 peptide; and *E. coli* zupT_OM + A7 peptide. Cultures were grown to an OD₆₀₀ of 0.2–0.3, at which point expression of particle synthesis pathways were induced with IPTG, arabinose, and/or anhydrotetracycline (aTc), as appropriate for each strain. Following induction, cultures were incubated for 2 h at 37 °C in the presence of 100 µM sodium thiosulfate to allow H₂S production. Cells were harvested by centrifugation and washed three times with Tris-HCl buffer (pH 7.0). Cadmium chloride was subsequently added to a final concentration of 1 µM, and the cultures were incubated for an additional 2 h at 37 °C without shaking.

### Characterization of CdS Quantum Dots

Absorbance and fluorescence emission spectra of the biosynthesized CdS quantum dots were measured using a TECAN Infinite M200 Pro plate reader, with excitation at 365 nm for fluorescence measurements. The size distribution of the quantum dots was determined via dynamic light scattering (DLS) using a Wyatt ZetaStar/Mobius instrument, and the hydrodynamic diameter was recorded.

## Acknowledgements

We would like to thank Manasi Gangan for help with the early stages of this project and the USC Nanobiophysics Core for assistance with dynamic light scattering measurements. We thank Matthew Lux, Greg Ellis, and Meghna Thakur for providing the pCad1 plasmid. This project was supported by Army Research Office under cooperative agreement number W911NF-24-2-0003 and by Office of Naval Research grant N00014-18-1-2632.

## References

Alivisatos, A. P. (1996). Semiconductor clusters, nanocrystals, and quantum dots. Science, 271(5251), 933–937.

Baba, T., Ara, T., Hasegawa, M., Takai, Y., Okumura, Y., Baba, M., Datsenko, K. A., Tomita, M., Wanner, B. L., & Mori, H. (2006). Construction of Escherichia coli K-12 in-frame, single-gene knockout mutants: the Keio collection. Molecular systems biology, 2, 2006.0008.

Bai, H., Zhang, Z., Guo, Y., & Jia, W. (2009). Biological synthesis of size-controlled cadmium sulfide nanoparticles using immobilized Rhodobacter sphaeroides. Nanoscale research letters, 4(7), 717.

Beabout, K., Bernhards, C. B., Thakur, M., Turner, K. B., Cole, S. D., Walper, S. A., … & Lux, M. W. (2021). Optimization of heavy metal sensors based on transcription factors and cell-free expression systems. ACS Synthetic Biology, 10(11), 3040–3054.

Bruna, N., Collao, B., Tello, A., Caravantes, P., Díaz-Silva, N., Monrás, J. P., … & Pérez-Donoso, J. M. (2019). Synthesis of salt-stable fluorescent nanoparticles (quantum dots) by polyextremophile halophilic bacteria. Scientific reports, 9(1), 1953.

Campoy, S., Jara, M., Busquets, N., Pérez de Rozas, A. M., Badiola, I., & Barbé, J. (2002). Role of the high-affinity zinc uptake znuABC system in Salmonella enterica serovar typhimurium virulence. Infection and immunity, 70(8), 4721–4725.

Chellamuthu, P., Tran, F., Silva, K. P. T., Chavez, M. S., El-Naggar, M. Y., & Boedicker, J. Q. (2019). Engineering bacteria for biogenic synthesis of chalcogenide nanomaterials. Microbial Biotechnology, 12(1), 161–172.

Chen, G., Yi, B., Zeng, G., Niu, Q., Yan, M., Chen, A., … & Zhang, Q. (2014). Facile green extracellular biosynthesis of CdS quantum dots by white rot fungus *Phanerochaete chrysosporium*. Colloids and Surfaces B: Biointerfaces, 117, 199–205.

Chen, Y. L., Tuan, H. Y., Tien, C. W., Lo, W. H., Liang, H. C., & Hu, Y. C. (2009). Augmented biosynthesis of cadmium sulfide nanoparticles by genetically engineered *Escherichia coli*. Biotechnology progress, 25(5), 1260–1266.

Contreras, F., Rivero, K., Rivas-Pardo, J. A., Liendo, F., Segura, R., Neira, N., Arenas-Salinas, M., Cortez-San Martín, M., & Arenas, F. (2025). Biosynthesis of Gold Nanostructures and Their Virucidal Activity Against Influenza A Virus. International journal of molecular sciences, 26(5), 1934.

Croce, A. C., & Bottiroli, G. (2014). Autofluorescence spectroscopy and imaging: a tool for biomedical research and diagnosis. European journal of histochemistry: EJH, 58(4), 2461.

Darroudi, M., Ahmad, M. B., Zamiri, R., Zak, A. K., Abdullah, A. H., & Ibrahim, N. A. (2011). Time-dependent effect in green synthesis of silver nanoparticles. International journal of nanomedicine, 677–681.

Dundas, C. M., Graham, A. J., Romanovicz, D. K., & Keitz, B. K. (2018). Extracellular Electron Transfer by *Shewanella oneidensis* Controls Palladium Nanoparticle Phenotype. ACS synthetic biology, 7(12), 2726–2736.

Edmundson, M. C., Capeness, M., & Horsfall, L. (2014). Exploring the potential of metallic nanoparticles within synthetic biology. New biotechnology, 31(6), 572–578.

Endo, G., & Silver, S. (1995). CadC, the transcriptional regulatory protein of the cadmium resistance system of *Staphylococcus aureus* plasmid pI258. Journal of bacteriology, 177(15), 4437–4441.

Era, Y., Dennis, J.A., Horsfall L.E., and Wallace, S. (2022). Palladium Nanoparticles from *Desulfovibrio alaskensis* G20 catalyze biocompatible sonogashira and biohydrogenation cascades, JACS Au 2022 2 (11), 2446–2452.

Gangan, M. S., Naughton, K. L., & Boedicker, J. Q. (2023). Utilizing a divalent metal ion transporter to control biogenic nanoparticle synthesis. Journal of Industrial Microbiology and Biotechnology, 50(1), kuad020.

Gu, X., Li, X., Zhang, R., Zheng, R., Li, M., Huang, R., & Pang, X. (2025). Isolation and characterization of a novel highly efficient bacterium Lysinibacillus boronitolerans QD4 for quantum dot biosynthesis. Frontiers in microbiology, 16, 1521632.

Houtepen, A. J., Sargent, E. H., Infante, I., Owen, J. S., Green, P. B., Schaller, R. D., … & Hens, Z. (2025). Colloidal quantum dots for optoelectronics. Nature Reviews Methods Primers, 5(1), 42.

Jia, Q. Y., Jia, R., Chen, C. M., & Wang, L. (2023). Characterization of CdSe QDs biosynthesized by a recombinant Rhodopseudomonas palustris. Biochemical Engineering Journal, 191, 108771.

Jiang, Y., Chen, B., Duan, C., Sun, B., Yang, J., & Yang, S. (2015). Multigene editing in the *Escherichia coli* genome via the CRISPR-Cas9 system. Applied and environmental microbiology, 81(7), 2506–2514.

Kang, S. H., Bozhilov, K. N., Myung, N. V., Mulchandani, A., & Chen, W. (2008). Microbial synthesis of CdS nanocrystals in genetically engineered *E. coli*. Angewandte Chemie International Edition, 47(28), 5186–5189.

Lakowicz, J. R., Gryczynski, I., Gryczynski, Z., & Murphy, C. J. (1999). Luminescence spectral properties of CdS nanoparticles. The Journal of Physical Chemistry B, 103(36), 7613–7620.

Le, N., & Kim, K. (2023). Current advances in the biomedical applications of quantum dots: promises and challenges. International Journal of Molecular Sciences, 24(16), 12682.

Lei, W., Liu, J., Liu, Y., Xu, J., & Wang, W. (2025). Overexpression of Tetrahymena cysteine synthetase 1 promotes cadmium removal by biosynthesizing cadmium sulfide quantum dots in Escherichia coli. International Journal of Molecular Sciences, 26(8), 3685.

Li, X. J., Wang, T. Q., Qi, L., Li, F. W., Xia, Y. Z., Zhang, C. J., … & Lin, J. Q. (2025). A one-step route for the conversion of Cd waste into CdS quantum dots by Acidithiobacillus sp. via unique biosynthesis pathways. RSC Chemical Biology, 6(2), 281–294.

Liu, J., Zheng, D., Zhong, L., Gong, A., Wu, S., & Xie, Z. (2021). Biosynthesis of biocompatibility Ag2Se quantum dots in *Saccharomyces cerevisiae* and its application. Biochemical and Biophysical Research Communications, 544, 60–64.

Liu, R., Shan, Y., Xi, S., Zhang, X., & Sun, C. (2022). A deep-sea sulfate-reducing bacterium generates zero-valent sulfur via metabolizing thiosulfate. Mlife, 1(3), 257–271.

Mao, C., Solis, D. J., Reiss, B. D., Kottmann, S. T., Sweeney, R. Y., Hayhurst, A., … & Belcher, A. M. (2004). Virus-based toolkit for the directed synthesis of magnetic and semiconducting nanowires. Science, 303(5655), 213–217.

Mi, C., Wang, Y., Zhang, J., Huang, H., Xu, L., Wang, S., … & Xu, S. (2011). Biosynthesis and characterization of CdS quantum dots in genetically engineered *Escherichia coli*. Journal of biotechnology, 153(3-4), 125–132.

Michalet, X., Pinaud, F. F., Bentolila, L. A., Tsay, J. M., Doose, S. J. J. L., Li, J. J., … & Weiss, S. (2005). Quantum dots for live cells, in vivo imaging, and diagnostics. science, 307(5709), 538–544.

Monrás, J. P., Díaz, V., Bravo, D., Montes, R. A., Chasteen, T. G., Osorio-Román, I. O., … & Pérez-Donoso, J. M. (2012). Enhanced glutathione content allows the in vivo synthesis of fluorescent CdTe nanoparticles by Escherichia coli. PLoS One, 7(11), e48657.

Narayanan, K. B., & Sakthivel, N. (2010). Biological synthesis of metal nanoparticles by microbes. Advances in colloid and interface science, 156(1-2), 1–13.

Naughton, K. L., & Boedicker, J. Q. (2021). Simulations to aid in the design of microbes for synthesis of metallic nanomaterials. ACS Synthetic Biology, 10(12), 3475–3488.

Órdenes-Aenishanslins, N. A., Saona, L. A., Durán-Toro, V. M., Monrás, J. P., Bravo, D. M., & Pérez-Donoso, J. M. (2014). Use of titanium dioxide nanoparticles biosynthesized by Bacillus mycoides in quantum dot sensitized solar cells. Microbial cell factories, 13(1), 90.

Park, Y., Espah Borujeni, A., Gorochowski, T.E. et al. Precision design of stable genetic circuits carried in highly-insulated *E. coli* genomic landing pads. Mol Syst Biol 16, MSB209584 (2020). 10.15252/msb.20209584

Patzer, S. I., & Hantke, K. (1998). The ZnuABC high-affinity zinc uptake system and its regulator Zur in Escherichia coli. Molecular microbiology, 28(6), 1199–1210.

Peng, X., Manna, L., Yang, W., Wickham, J., Scher, E., Kadavanich, A., & Alivisatos, A. P. (2000). Shape control of CdSe nanocrystals. Nature, 404(6773), 59–61.

Peng, Z.A., & Peng, X. (2001). Formation of high-quality CdTe, CdSe, and CdS nanocrystals using CdO as precursor. Journal of the American Chemical Society, 123(1), 183–184.

Qu, L., & Peng, X. (2002). Control of photoluminescence properties of CdSe nanocrystals in growth. Journal of the American Chemical Society, 124(9), 2049–2055.

Reisch, C. R., & Prather, K. L. (2015). The no-SCAR (Scarless Cas9 Assisted Recombineering) system for genome editing in *Escherichia coli*. Scientific reports, 5(1), 15096.

Shafiei, H., Samadi-Maybodi, A., & Mohseni, M. (2025). Optical properties of novel synthesized ZrS2 quantum dots based on Escherichia coli bacteria. Spectrochimica Acta Part A: Molecular and Biomolecular Spectroscopy, 337, 126113.

Shatalin, K., Shatalina, E., Mironov, A., & Nudler, E. (2011). H2S: a universal defense against antibiotics in bacteria. Science, 334(6058), 986–990.

Sharma, D., Kanchi, S., & Bisetty, K. (2019). Biogenic synthesis of nanoparticles: a review. Arabian journal of chemistry, 12(8), 3576–3600.

Shivaji, K., Mani, S., Ponmurugan, P., De Castro, C. S., Lloyd Davies, M., Balasubramanian, M. G., & Pitchaimuthu, S. (2018). Green-synthesis-derived CdS quantum dots using tea leaf extract: antimicrobial, bioimaging, and therapeutic applications in lung cancer cells. ACS Applied Nano Materials, 1(4), 1683–1693.

Siegel, L. M. (1965). A direct microdetermination for sulfide. Analytical biochemistry, 11(1), 126–132.

Syed, A., & Ahmad, A. (2013). Extracellular biosynthesis of CdTe quantum dots by the fungus *Fusarium oxysporum* and their anti-bacterial activity. Spectrochimica Acta Part A: Molecular and Biomolecular Spectroscopy, 106, 41–47.

Taudte, N., & Grass, G. (2010). Point mutations change specificity and kinetics of metal uptake by ZupT from *Escherichia coli*. Biometals, 23(4), 643–656.

Vargas-Reyes, M., Bruna, N., Ramos-Zúñiga, J., Valenzuela-Ibaceta, F., Rivas-Álvarez, P., Navarro, C. A., & Pérez-Donoso, J. M. (2024). Biosynthesis of photostable CdS quantum dots by UV-resistant psychrotolerant bacteria isolated from Union Glacier, Antarctica. Microbial Cell Factories, 23(1), 140.

Vo, P. L. H., Ronda, C., Klompe, S. E., Chen, E. E., Acree, C., Wang, H. H., & Sternberg, S. H. (2021). CRISPR RNA-guided integrases for high-efficiency, multiplexed bacterial genome engineering. Nature biotechnology, 39(4), 480–489.

Yan, Z. Y., Du, Q. Q., Qian, J., Wan, D. Y., & Wu, S. M. (2017). Eco-friendly intracellular biosynthesis of CdS quantum dots without changing Escherichia coli’s antibiotic resistance. Enzyme and Microbial Technology, 96, 96–102.

Yu, W. W., Wang, Y. A., & Peng, X. (2003). Formation and stability of size-, shape-, and structure-controlled CdTe nanocrystals: ligand effects on monomers and nanocrystals. Chemistry of Materials, 15(22), 4300–4308.

Zhang, Z., Chen, J., Yang, Q., Lan, K., Yan, Z., & Chen, J. (2018). Eco-friendly intracellular microalgae synthesis of fluorescent CdSe QDs as a sensitive nanoprobe for determination of imatinib. Sensors and Actuators B: Chemical, 263, 625–633.

Zhou, Z., Pu, Y., Liu, W., Jin, M., & Xian, M. (2022). Assembly and synthesis mechanism of CdSe quantum dots in recombinant *Escherichia coli* expressing metallothionein. ACS Sustainable Chemistry & Engineering, 11(1), 113–121.

Zhu, W., & Chu, W. (2022). A sensitive visual method for the detection of hydrogen sulfide producing bacteria. Journal of Visualized Experiments (JoVE), (184), e64201.

